# AMPK regulates germline stem cell integrity and quiescence through a *mir-1/tbc-7/rab-7* pathway in *C. elegans*

**DOI:** 10.1101/2021.09.22.461433

**Authors:** Christopher Wong, Pratik Kadekar, Elena Jurczak, Richard Roy

**Affiliations:** Department of Biology, McGill University, Montreal, Quebec, Canada

**Author notes:** Corresponding author (RR).

## Abstract

During periods of energetic stress, *Caenorhabditis elegans* can undergo a global quiescent stage known as “dauer”. During this stage, all germline stem cells undergo G2 cell cycle arrest through an AMPK-dependent mechanism. In animals that lack AMPK signalling, the germ cells fail to arrest, undergo uncontrolled proliferation and lose their reproductive capacity. These germline defects are accompanied by an altered chromatin landscape and gene expression program. We identified an allele of *tbc-7*, a RabGAP protein that functions in the neurons, which when compromised, suppresses the germline hyperplasia in the dauer larvae, as well as the post-dauer sterility and somatic defects characteristic of AMPK mutants. This mutation also corrects the abundance and aberrant distribution of transcriptionally activating and repressive chromatin marks in animals that otherwise lack all AMPK signalling. We identified RAB-7 as one of the potential RAB proteins that is regulated by *tbc-7* and show that the activity of RAB-7 is critical for the maintenance of germ cell integrity during the dauer stage. A singular small RNA, *mir-1*, was identified as a direct negative regulator of *tbc-7* expression through the analysis of seed sequences on the 3’UTR of *tbc-7*. Animals lacking *mir-1* are post-dauer sterile, displaying a similar phenotype to AMPK mutants. Altogether, our findings describe a novel *mir-1/tbc-7/rab-7* pathway occurring in the neurons that regulates the germ line in a cell non-autonomous manner.

## Introduction

For many organisms, development unravels as a successive series of growth, quiescence and differentiation phases that proceed according to regulations dictated by an organism-specific program. This is particularly true for closed systems, like during embryonic development, where events often unfold independent of external resource availability due to maternal provisioning for the developing organism.

This situation changes dramatically as animals emerge from the embryo, and juveniles are subject to dramatic fluctuations in growing conditions. These challenges can be detrimental to continuous development and thus have driven the emergence of diverse mechanisms of adaptation that enhance survival. The execution of a quiescent state as a means to circumvent these challenges is widespread. In microbial cells, like bacteria and yeast, rapid growth and divisions arrest when resources are exhausted, while cells remain viable and ready to resume proliferation when growth conditions improve [1–3].

Quiescence is by no means unique to microbes. Mammalian cells in culture arrest their cell divisions and enter a G_0_ state if they are deprived of growth factors (serum), glucose, or when they become confluent [4]. Moreover, cells that no longer appropriately execute quiescence in response to these cues often demonstrate other rogue behaviours, typical of transformed cells [2]. It is therefore highly advantageous to appropriately respond to these adverse environmental signals and execute a quiescent state as an adaptive means of protecting the cell.

Multicellular organisms also employ various states of quiescence to ensure survival during periods of environmental stress [5–7]. However, not all cells in the organism are capable of sensing these cues and therefore depend on specialized cells to communicate information to their distant neighbours to protect them against these challenges.

Many animals, including humans, forego reproduction during periods of excess stress or starvation [7, 8]. Both starved *C. elegans,* or mutants that lack insulin signalling are infertile and do not produce oocytes [9]. However, this oogenesis defect can be bypassed in these mutants by eliminating the stress-responsive sensor AMP-activated protein kinase (AMPK) [8, 10]. Activation of AMPK as animals execute the highly-resistant, diapause-like dauer stage results in germline quiescence, while other somatic cell divisions are arrested through a parallel pathway [10, 11]. In the absence of AMPK signalling, mutant germ cells continue to proliferate despite a lack of resources, resulting in highly penetrant sterility following recovery [8, 10].

The dauer stage can last for long periods and yet when animals exit this diapause, they are fully fertile with little or no negative reproductive consequence [12, 13]. On the other hand, animals that lack all AMPK signalling exhibit highly penetrant sterility after emerging from the dauer stage. The AMPK-mediated quiescence that occurs during the dauer stage is therefore protective and is critical to preserve germ cell integrity during this period of prolonged stress. Consistent with this, wild-type animals that transit through the dauer stage exhibit differences in their gene expression compared to animals that never encountered the stresses associated with dauer development [14, 15]. These changes persist in the post-dauer animals, as a molecular memory stored in the form of epigenetic marks, which have been shown to influence post-dauer fertility.

Mutants that lack all AMPK signalling display significant changes in gene expression during the dauer stage and as post-dauer adult animals when compared to wild type animals [10]. In addition, the levels of transcriptionally activating and repressive chromatin marks within the germ line are abnormally upregulated, while their distribution is also abnormal. This suggests that the differences in gene expression in AMPK mutants could be attributed to the misregulation of chromatin marks in the affected germ cells.

Recent studies have indicated that AMPK can act cell non-autonomously to regulate life span [8], while the restoration of AMPK in some somatic cells was sufficient to correct the germline defects in AMPK mutants [10, 16]. Moreover, when components of the endogenous small RNA pathway are compromised, the post-dauer sterility and the germline hyperplasia typical of AMPK mutants are also partially suppressed. These examples highlight how information is conveyed to the germ line through specialized sensing cells, very often by neurons. In the context of dauer development, AMPK may modulate an endogenous small RNA pathway in the soma, or perhaps even more specifically in the neurons, to regulate the development of the germ line in response to the metabolic challenges associated with dauer development.

In this study, we define a pathway in which AMPK regulates germ cell homeostasis through a small RNA-based mechanism that is engaged in the neurons. By performing a genetic screen designed to identify genes involved in the AMPK regulation of germ cell integrity and germline quiescence during the dauer stage, we isolated eight alleles that partially restore germ cell function in these mutants. One of these genes is a RabGAP that presumably enhances the inherent GTP-hydrolysis ability of one or more Rab proteins, converting it from its active RAB-GTP form to its inactivated RAB-GDP form. We show that this RabGAP functions in the neurons to negatively regulate its RAB target. Furthermore, we demonstrate that the RabGAP is subject to regulation by a single microRNA, thus defining a small RNA-based pathway in the neurons that is critical for the maintenance of germ cell integrity during this period of developmental quiescence.

## Results

### Mutations that reverse the germline defects of AMPK mutants

The *C. elegans* dauer stage is marked by a pervasive organismal developmental quiescence that is initiated as animals transit through the preparatory L2d stage. This is accompanied by a complete arrest in germ cell divisions mediated by the activity of the *C. elegans* orthologues of the human tumour suppressor gene LKB1 (*par-4*) and one of its many downstream targets, AMPK [8]. Animals that lack all AMPK signalling (*aak(0)*) undergo germline hyperplasia during dauer formation and exhibit post-dauer sterility, in addition to a number of metabolic/somatic phenotypes [10]. Curiously, the compromise of various components involved in the biogenesis and function of small RNAs can partially restore germ cell quiescence in AMPK mutant dauer larvae, in addition to the corresponding post-dauer fertility in these mutants.

To identify additional genes that converge either on these small RNA regulators or on AMPK targets in the germ line, we performed a forward genetic screen to obtain mutants that suppress the germline defects typical of AMPK mutants. Eight alleles were isolated that fall into five complementation groups (Table 1), each of which was able to suppress the germ cell hyperplasia in the AMPK mutant dauer larvae, and/or partially restore post-dauer fertility in AMPK mutants (Fig S1A-C and Table S1).

**Table 1.** Mutants that suppress the sterility of post-dauer AMPK mutants fall into five complementation groups.

| Allele(s) | Complementation Group | Dominance/ Recessiveness | Penetrance |  |  |
| --- | --- | --- | --- | --- | --- |
|  |  |  | % egg laying animals | No. of germ cells | % wild type-like |
| Control | - | - | 100 | 35 | 100 |
| *aak(0)* | - | - | 0 | 116 | 68 |
| *rr166* | Group 1 | Recessive | 91 | 68 | 92 |
| *rr249* | Group 2 | Semi-dominant | 78 | 97 | 84 |
| *rr256* | Group 3 | Recessive | 67 | 85 | 84 |
| *rr261* | Group 4 | Recessive | 67 | 90 | 76 |
| *rr266* | Group 3 | Recessive | 63 | 66 | 84 |
| *rr267* | Group 1 | Recessive | 67 | 97 | 80 |
| *rr268* | Group 5 | Recessive | 68 | 75 | 76 |
| *rr289* | Group 3 | Recessive | 61 | 112 | 72 |

Eight EMS-generated alleles were isolated from a genetic screen designed to identify genes involved in the AMPK regulation of germ cell integrity and germline quiescence during the dauer stage. Through cross complementation analysis they were found to fall into five complementation groups, two of which had multiple independent alleles; Group 1 - *rr166, rr267* and Group 3 - *rr256, rr266, rr289.* Based on our F_1_ and F_2_ cross-complementation analysis all the alleles behave recessively, with the exception of *rr249*, which acts semi-dominantly.

Our further analysis indicated that all the isolated alleles are recessive with the exception of *rr249*, which is semi-dominant (Table 1). One of these alleles, *rr166,* was able to suppress the AMPK germline defects to a greater extent when compared to others (Fig 1A-C). Moreover, *rr166* also suppressed most of the post-dauer somatic defects, suggesting that this allele may signal not only the germ line but also the somatic tissues to adjust cell division timing for the duration of the diapause (Table S1).

**Fig 1.**
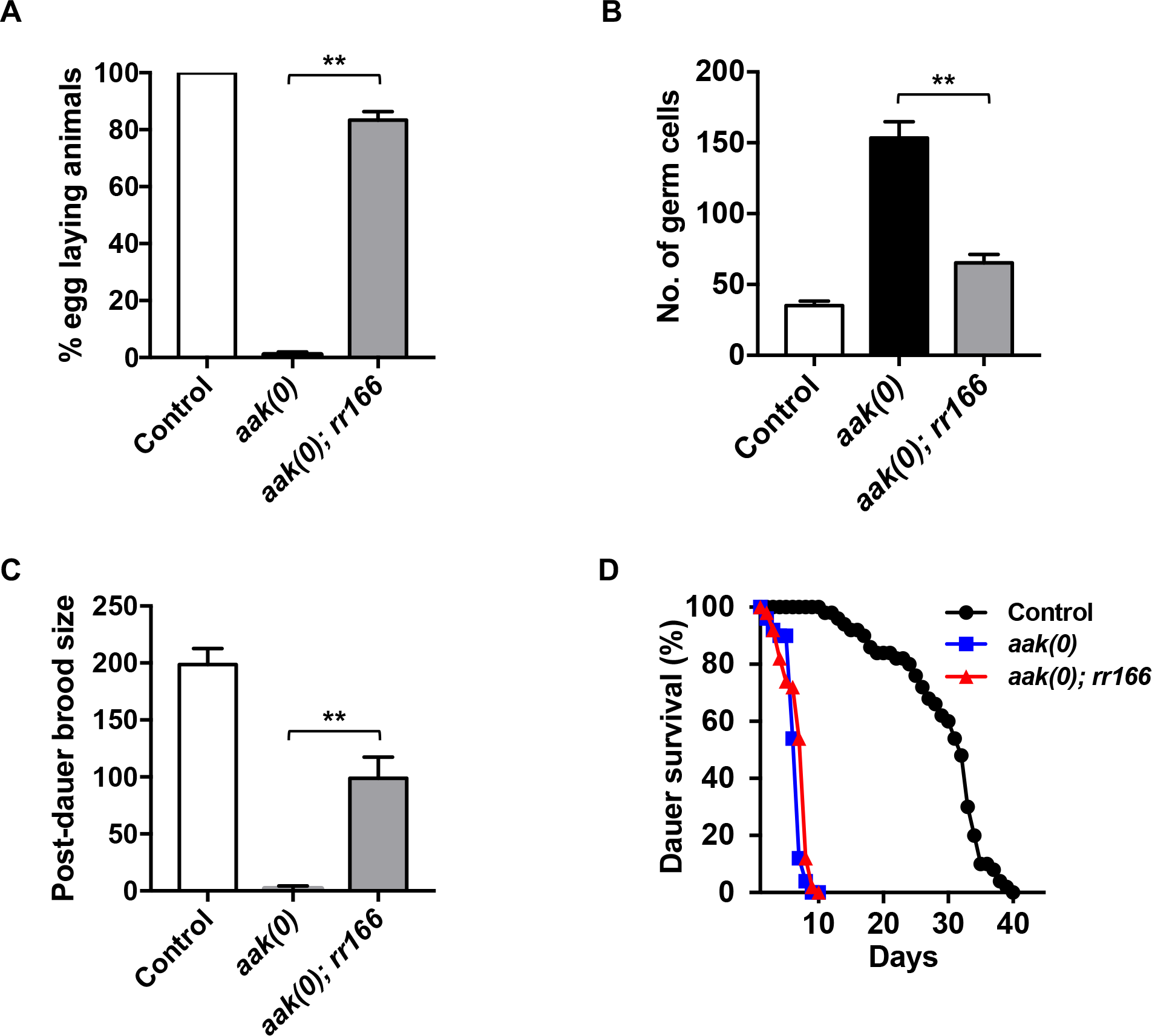
*rr166* partially suppresses the post-dauer sterility and dauer germline hyperplasia of AMPK mutants. (A-C) *rr166* suppresses the (A) post-dauer sterility, (B) dauer germline hyperplasia, and (C) brood size defects in animals that lack all AMPK signalling. (D) *rr166* does not suppress the reduced dauer survival typical of *aak(0)* mutants. ***P < 0.0001 when compared to *aak(0)* using ordinary one-way ANOVA for post-dauer brood size and no. of germ cells. Data are represented as mean ± SD. ***P < 0.0001 when compared to *aak(0)* using Marascuilo procedure for % egg laying animals (% total). No significant difference (ns) was observed when compared to *aak(0)* using log-rank test for dauer survival. All animals carry the *daf-2(e1370)* allele. n = 50.

AMPK is required for the typical long-term survival of the dauer larvae by regulating triglyceride hydrolysis during the dauer stage [16]. Dauer larvae with intact AMPK signalling can survive for several weeks, while mutants with disrupted AMPK/LKB1 signalling die prematurely due to the inappropriate depletion of stored lipids important to sustain animals for the duration of the diapause. Strikingly, *rr166* did not improve the dauer survival of *aak(0)* mutants, suggesting that the *rr166* mutation does not affect other AMPK-regulated pathways and may be specific to germ line/gonadal physiology (Fig 1D).

### *rr166* corrects the abnormal deposition of the chromatin marks in the germ line of AMPK mutant dauer larvae and post-dauer adults

*aak(0)* mutants not only have upregulated levels of both transcriptionally activating and repressive chromatin marks, but they also have an abnormal distribution of these epigenetic modifications across the germ line in both dauer and post-dauer animals [10]. This altered chromatin landscape is associated with a dramatic change in germline gene expression during the dauer stage which remains unaltered upon dauer exit, persisting into the post-dauer adult. To better understand how *rr166* suppresses the AMPK germline defects, we examined the levels of chromatin marks that were aberrant in the AMPK mutant dauer larvae to assess how these epigenetic modifications are affected in the *rr166* mutant. Western analysis revealed that both activating (H3K4me3) and repressive (H3K9me3) chromatin modifications were restored to nearly normal levels in the germ cell nuclei of *rr166* dauer larvae that lack all AMPK signalling (Fig 2A-A’’). In addition, the distribution of the chromatin marks was also corrected (Fig 2B-C’’).

**Fig 2.**
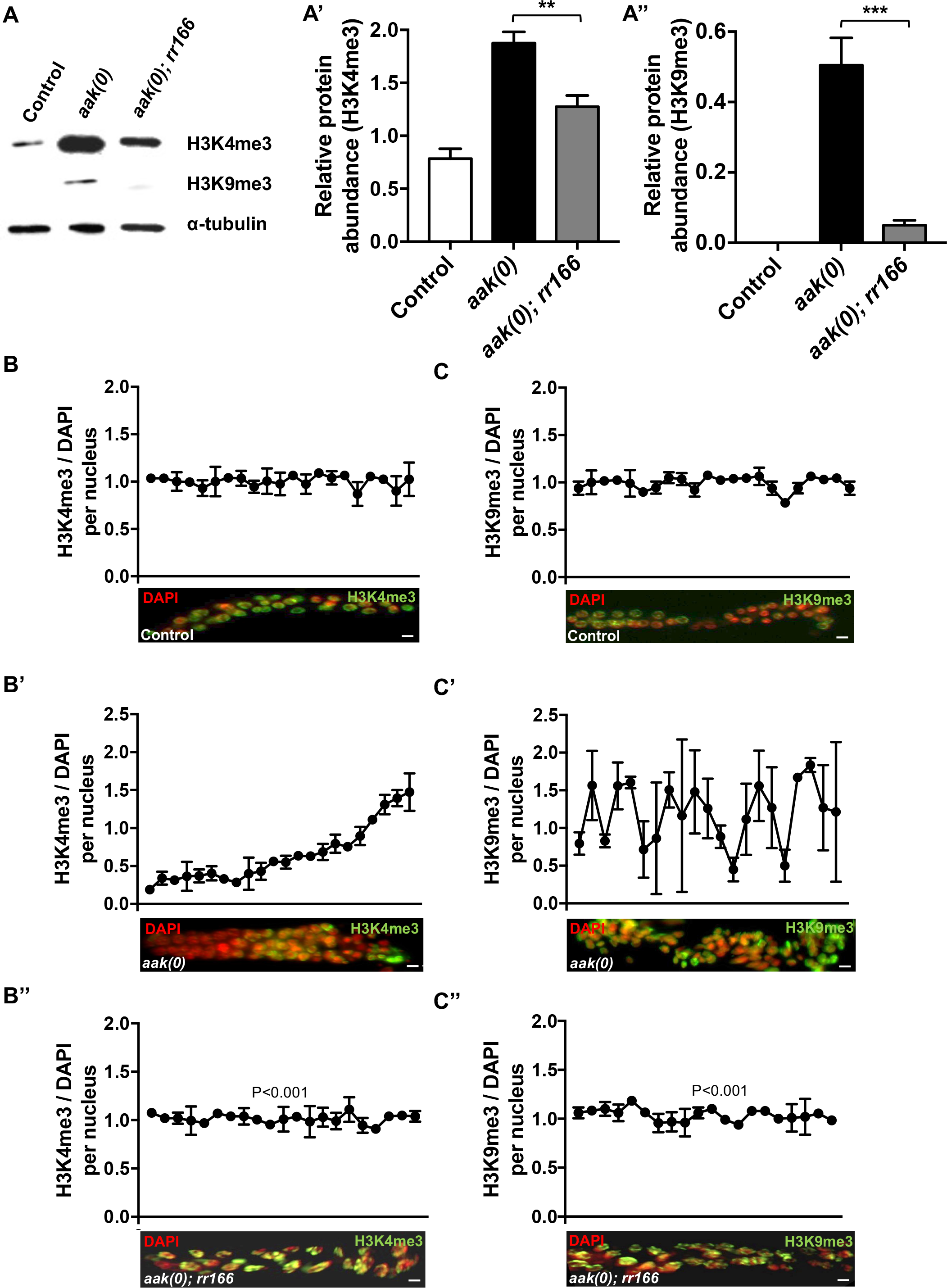
The abnormal abundance and distribution of chromatin marks typical of dauer larvae that lack AMPK is corrected in *rr166* mutants. (A) Global levels of H3K4me3 and H3K9me3 were quantified by performing whole-animal western analysis on dauer larvae. (A’-A’’) Levels of chromatin marks were quantified and normalized to α-tubulin using ImageJ software. ***P < 0.0001 when compared to *aak(0)* using Student’s *t* test. (B-C’’) The distribution and abundance of activating and repressive chromatin marks are corrected in the *rr166* mutants. The top row (Control), middle row (*aak(0)*), and bottom row (*aak(0); rr166*) show (B, B’, B’’) H3K4me3 (green), (C, C’, C’’) H3K9me3 (green), and DAPI (red). The graphs represent the average immunofluorescence signal of anti-H3K4me3 and anti-H3K9me3 normalized to DAPI in each nucleus across the dissected germ line. All images are merged, condensed Z stacks and are aligned such that distal is left and proximal is right. Due to technical difficulties, only single gonadal arms were analyzed (distal, proximal). **P < 0.001 using the F-test for variance when compared to *aak(0)*. All animals carry the *daf-2(e1370)* allele. Scale bar: 4 μm. n = 15.

Using the same approach, we confirmed that the *rr166* mutation was also able to suppress the abnormal upregulation of the chromatin marks in the *aak(0)* mutants during the post-dauer stage as well (Fig 3A-A’’). Immunofluorescence analysis against these histone modifications indicates that *rr166* can correct the abundance of each mark in individual nuclei, while also reestablishing the overall pattern across the germ line (Fig 3B-C’’). Interestingly, while the levels of H3K9me3 in the *rr166* mutants are similar compared to control animals, the levels of H3K4me3 are not completely restored to control levels, albeit reduced compared to those of *aak(0)* mutants (Fig 3B-B’’). Together, these data indicate that *rr166* corrects the abundance and distribution of chromatin marks in the germ line that presumably primes germline gene expression for a period of developmental quiescence in animals that lack all AMPK signalling.

**Fig 3.**
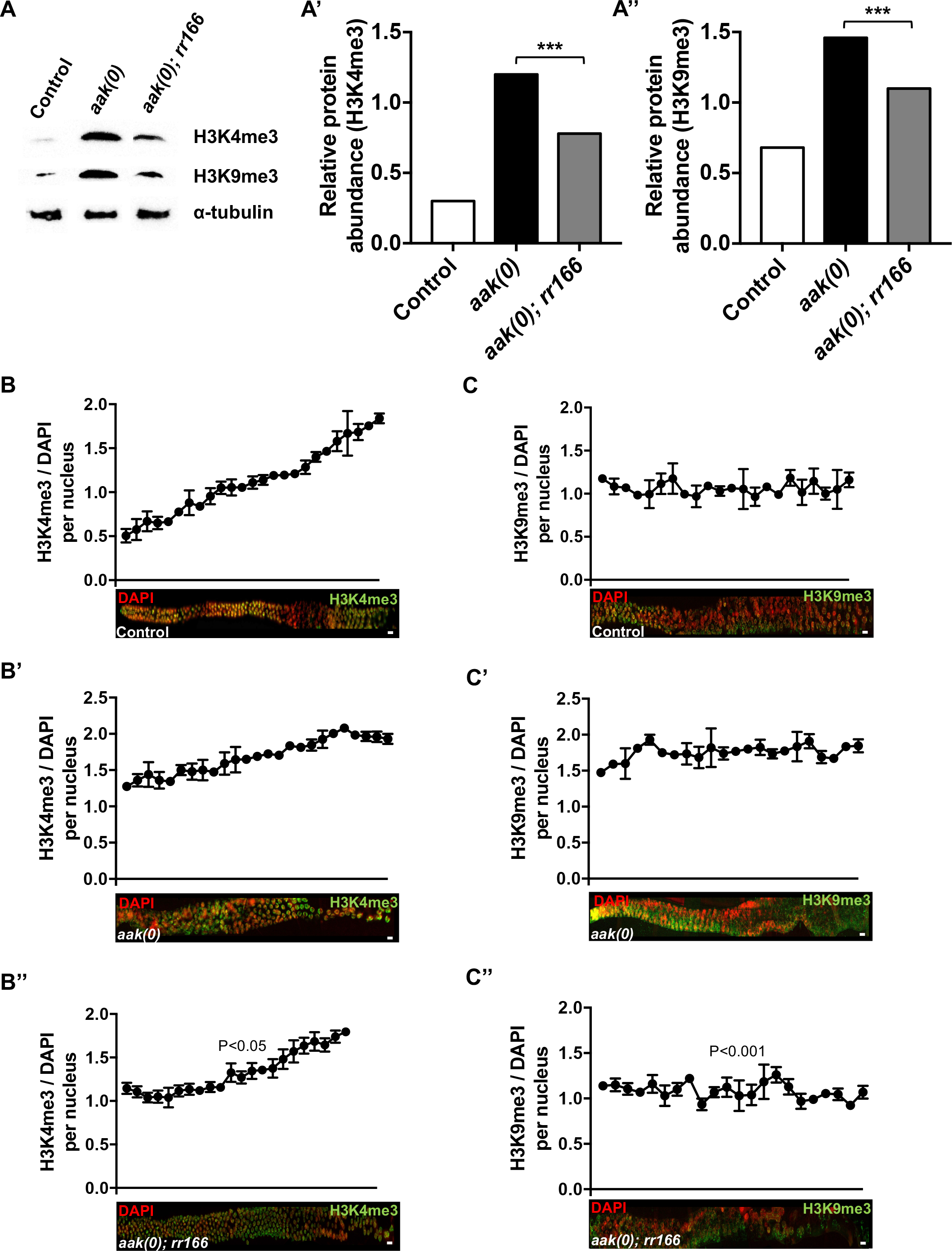
The *rr166* mutation restores appropriate chromatin remodeling in post-dauer AMPK mutants. (A) Global levels of H3K4me3 and H3K9me3 were quantified by performing whole-animal western analysis of dauer larvae. (B) Chromatin marks were quantified and normalized to α-tubulin using ImageJ software. ***P < 0.0001 when compared to *aak(0)* using Student’s *t* test. (B-C’’) The aberrant distribution and abundance of activating (H3K4me3) and repressive (H3K9me3) chromatin marks observed in post-dauer adults that lack AMPK signalling are corrected in the *rr166* mutants. The top row (Control), middle row (*aak(0)*), and bottom row (*aak(0); rr166*) show (B, B’, B’’) H3K4me3 (green), (C, C’, C’’) H3K9me3 (green), and DAPI (red). The graphs represent the average immunofluorescence signal of anti-H3K4me3 and anti-H3K9me3 normalized to DAPI across the dissected germ line. All images are merged, condensed Z stacks that are aligned such that distal is left and proximal is right. Due to technical difficulties, only single gonadal arms were analyzed (distal, proximal). **P < 0.001 using the F-test for variance when compared to *aak(0)*. All animals carry the *daf-2(e1370)* allele. Scale bar: 4 μm. n = 15.

### Misregulated RabGAP activity in neurons of AMPK mutants results in germ line abnormalities during the dauer stage

Genomic DNA obtained from *rr166* mutants was subjected to whole genome sequencing to identify relevant polymorphisms that could correspond to the affected gene responsible for the suppression of AMPK germline defects [17]. *rr166* was revealed to be a typical EMS-generated G->A transition mutation in a predicted RabGAP protein called *tbc-7*.

To confirm that *rr166* is indeed an allele of *tbc-7*, we performed RNAi against *tbc-7* in the *aak(0)* mutants. *aak(0)* mutants subjected to *tbc-7* RNAi showed no significant difference in their post-dauer fertility when compared to *aak(0); rr166* animals (Fig 4A). Furthermore, when *aak(0); rr166* mutants were injected with a wild type copy of *tbc-7*, the suppression of AMPK mutant germline defects was reverted, suggesting that *rr166* is an allele of *tbc-7* (Fig 4A). To determine if the *tbc-7(rr166)* point mutation is a null allele, we crossed the *tbc-7(rr166)* point mutation mutant with a mutant that contains a deletion of the entire *tbc-7* gene (*tm10766*). A homozygous *tm10766* mutation is lethal, therefore these mutants are balanced and maintained as heterozygotes. When *tm10766* hermaphrodites were crossed with *tbc-7(rr166)* males, we were able to isolate F1 mutants carrying the *tm10766* deletion and the *tbc-7(rr166)* point mutation, suggesting that *rr166* is a hypomorphic allele of *tbc-7* (Fig S2A-B).

**Fig 4.**
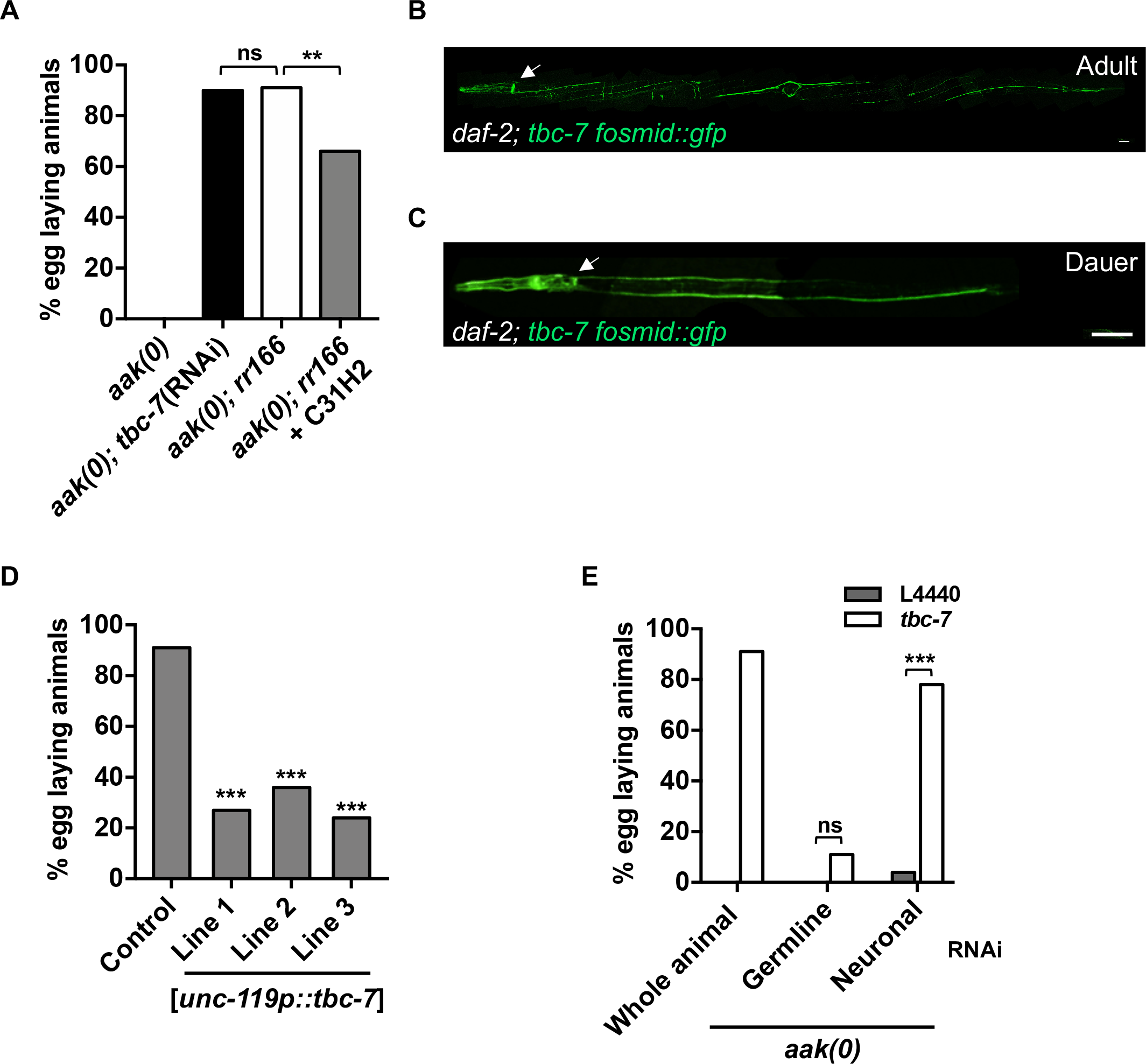
*tbc-7* functions in the neurons to maintain germ cell integrity during the dauer stage. (A) *rr166* was confirmed to be an allele of *tbc-7.* RNAi against *tbc-7* in *aak(0)* suppresses the post-dauer sterility. Injection of a cosmid containing a wild-type copy of *tbc-7* (C31H2) into *aak(0); tbc-7* mutants partially reverts the suppression of *aak(0)* germline defects. (B-C) The expression of a fosmid containing *tbc-7* translationally fused to GFP is shown in (B) wild-type adult animals and in (C) dauer larvae. White arrows indicate the nerve ring. Scale bars: 25 μm. Note that the morphological abnormality is a result of the *rol-6D* co-transformation marker. (D) A wild-type copy of *tbc-7* expressed exclusively in the neurons by the *unc-119* promoter in the *tbc-7*-suppressed mutants reverts the suppression of the AMPK germline phenotypes. (E) Tissue-specific RNAi experiments reveal that the compromise of *tbc-7* expression exclusively in the neurons (Neuronal) is sufficient to suppress the AMPK germline defects. ***P < 0.0001 when compared to controls based on Marascuilo procedure for % egg laying animals. All animals carry the *daf-2(e1370)* allele. n = 50.

Recent work demonstrated that AMPK regulates germ cell integrity cell non-autonomously during the dauer stage [10, 16]. Restoring AMPK function in the neurons and in the excretory system re-establishes germline quiescence and germ cell integrity in AMPK mutants [10]. Furthermore, the somatic expression of *aak-2* is able to correct both the abundance and distribution of chromatin marks in the dauer germ line, suggesting that the somatic function of AMPK controls the execution of germline quiescence in response to dauer cues.

*tbc-7* encodes a RabGAP protein that is conserved in *Drosophila* and in humans, where it is critical for appropriate synaptic vesicle dynamics in the neurons [18, 19]. Because of this neuronal role, we sought to determine if it also regulates the germ line non-autonomously during the dauer stage. To better understand where *tbc-7* is expressed we generated a transgenic strain using a fosmid containing a TBC-7::GFP translational fusion protein. Imaging revealed that TBC-7 is highly expressed in the neurons of adult animals grown in replete conditions as well as in the dauer larvae (Fig 4B-C). The neuronal expression of *tbc-7* is critical for its function, since the introduction of a wild-type copy of *tbc-7* driven by a pan-neuronal promoter in the *aak(0); tbc-7* mutants reverts the suppression of AMPK germline defects (Fig 4D).

To further confirm that *tbc-7* acts in the neurons, we used a tissue-specific RNAi strategy that allowed us to compromise the expression of *tbc-7* exclusively in one tissue at a time (see Materials and Methods) (Table S2) [20, 21]. If *tbc-7* activity is required in the neurons, then *aak(0)* mutants that are subjected to *tbc-7* RNAi exclusively in the neurons should phenocopy the *tbc-7(rr166)* mutation and suppress the AMPK germline defects. Using the neuronal RNAi strain, we found that *tbc-7* RNAi indeed suppresses the post-dauer sterility of *aak(0)* mutants, while strains that had *tbc-7* compromised exclusively in the germ line showed no such suppression (Fig 4E). Taken together, these data indicate that TBC-7 functions cell non-autonomously in the neurons to modulate AMPK-dependent germline quiescence and germ cell integrity during the dauer stage.

### TBC-7 negatively regulates RAB-7 in the neurons to maintain germ cell integrity

TBC-7 is a RabGAP protein that enhances the inherent GTP hydrolysis activity of Rab GTPases, converting them from their active GTP-bound form into their inactive GDP-bound form. In AMPK mutants, the *tbc-7*-associated RAB(s) are in their inactive GDP-bound form due to misregulated TBC-7 activity, while in *aak(0); tbc-7* mutants, the *tbc-7*-associated RAB(s) are in their active GTP-bound form (Fig 5A). Thus, compromising the expression of the RAB(s) associated with *tbc-7* should revert the suppression conferred by the *tbc-7* mutation, causing a loss of germ cell integrity and post-dauer fertility. To identify which RAB protein is regulated by *tbc-7*, each individual *rab* gene in the *C. elegans* genome was compromised using a hypomorphic RNAi strategy [22]. This RNAi survey of RAB proteins revealed that TBC-7 may regulate a large number of RAB effectors (Fig 5B and Table S3). However, when *rab-7* was compromised in the *aak(0); tbc-7* mutants, the post-dauer fertility was greatly reduced to a level similar to that of post-dauer *aak(0)* mutants, suggesting that *rab-7* is likely required for the correct regulation of germline quiescence during the dauer stage (Fig 5B). To confirm that RAB-7 is directly regulated by TBC-7 in the neurons, we once again used our tissue-specific RNAi system [20, 21] to knock down *rab-7* activity exclusively in the neurons, which resulted in highly penetrant post-dauer sterility (Table S4), while its compromise in the germ line had no observable effects on the reproductive output of the post-dauer animal (Fig 5C). Together, our analyses indicate that TBC-7 negatively regulates RAB-7 in the neurons to regulate the germ line during the dauer stage.

**Fig 5.**
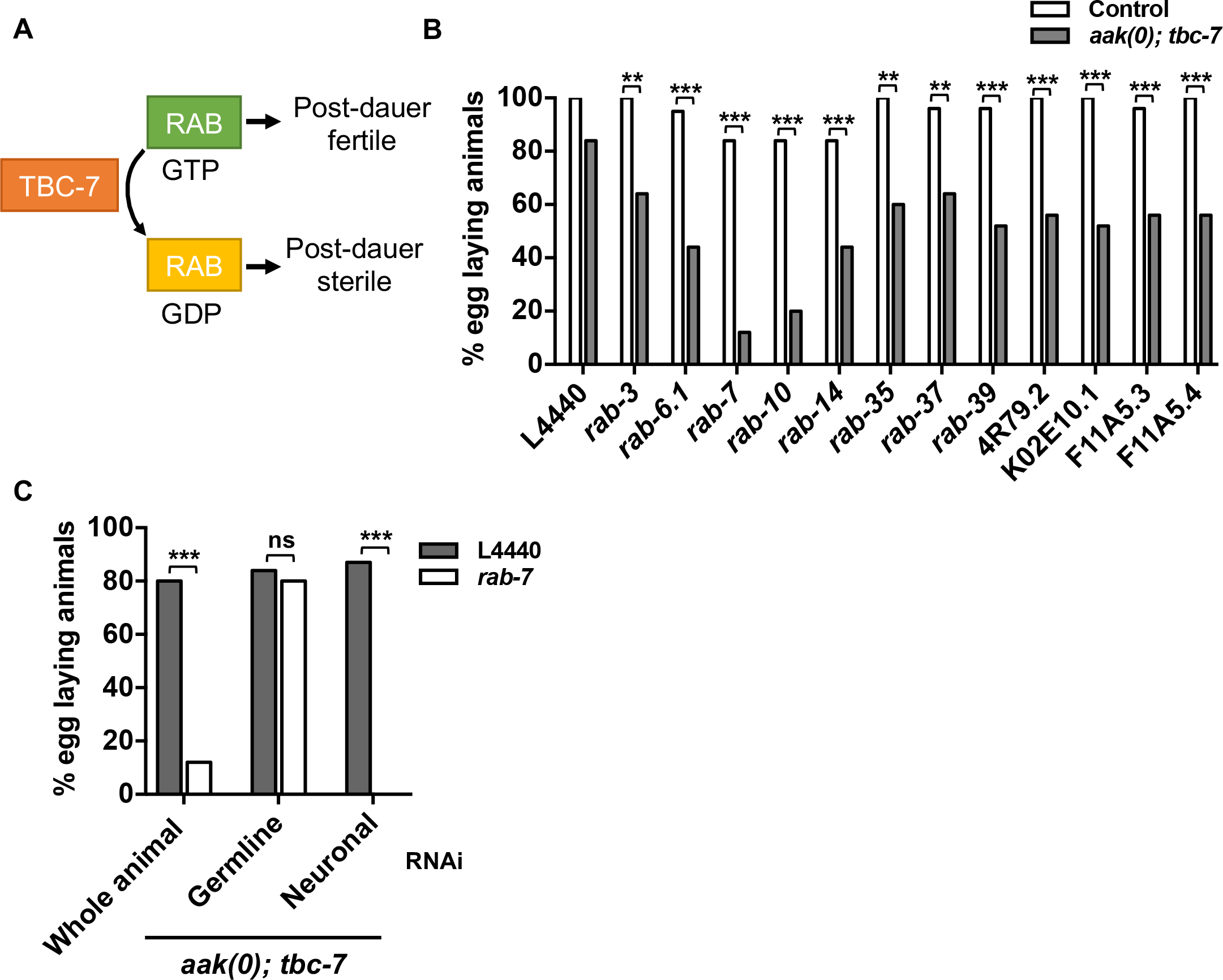
TBC-7 regulates RAB-7 in the neurons to maintain germ cell integrity during the dauer stage. (A) Graphical figure illustrating the regulation of RAB activity by TBC-7. In *aak(0); tbc-7* mutants, TBC-7 is inactive, allowing its RAB protein to be active in its GTP-bound state. By compromising the expression of this RAB protein using RNAi, the suppression of post-dauer sterility is lost. (B) By subjecting each predicted known *rab* gene in the *C. elegans* genome to RNAi, several RAB proteins were identified as potential RAB partners of TBC-7. ***P < 0.0001, **P < 0.001 when compared to control using Marascuilo procedure for % of egg laying animals. (C) Tissue-specific RNAi experiments reveal that *rab-7* expression is required in the neurons and not the germ line for the *tbc-7* suppression of AMPK germline phenotypes. ***P < 0.0001, **P < 0.001 when compared to L4440 empty vector using Marascuilo procedure for % of egg laying animals. All animals carry the *daf-2(e1370)* allele. n = 50.

### *mir-1* negatively regulates *tbc-7* downstream of AMPK activation to promote germline quiescence

Our previous findings suggest that small RNA homeostasis is misregulated in the germ line during the dauer stage in AMPK mutants since the compromise of the endogenous siRNA pathway partially suppresses the typical germline defects in these mutants [10]. Because the compromise of *tbc-7* is also capable of suppressing the germline defects of AMPK mutants, we hypothesized that *tbc-7* might also function through a small RNA pathway, or alternatively, be regulated itself by small RNAs. To determine if the suppression of the germline defects conferred by *tbc-7* requires the production of small RNAs, we performed RNAi on the *C. elegans* ortholog of Dicer (*dcr-1*), one of the critical upstream enzymes involved in small RNA biosynthesis [23]. The reduction of *dcr-1* in the *aak(0); tbc-7* mutants reverts the *tbc-7*-associated suppression of post-dauer sterility, suggesting that the production of small RNAs is required for this effect. To determine which class of small RNAs is required for the observed suppression, critical components that are required for the function of each DCR-1-dependent class of endogenous small RNAs (ALG-3/4-class 26G RNA, ERGO-1-class 26G RNA, and DRSH-1, PASH-1-microRNA) were disabled in the *aak(0); tbc-7* mutants [24–26]. When either of the *C. elegans* orthologs of Drosha and Pasha, *drsh-1* and *pash-1*, critical gene products required for microRNA biogenesis, were compromised in the *aak(0); tbc-7* mutants, the suppression of AMPK mutant germline defects was abrogated (Fig 6A). On the contrary, the disruption of various components involved in siRNA biogenesis had no effect on the post-dauer fertility of the *aak(0); tbc-7* mutants (Fig S3).

**Fig 6.**
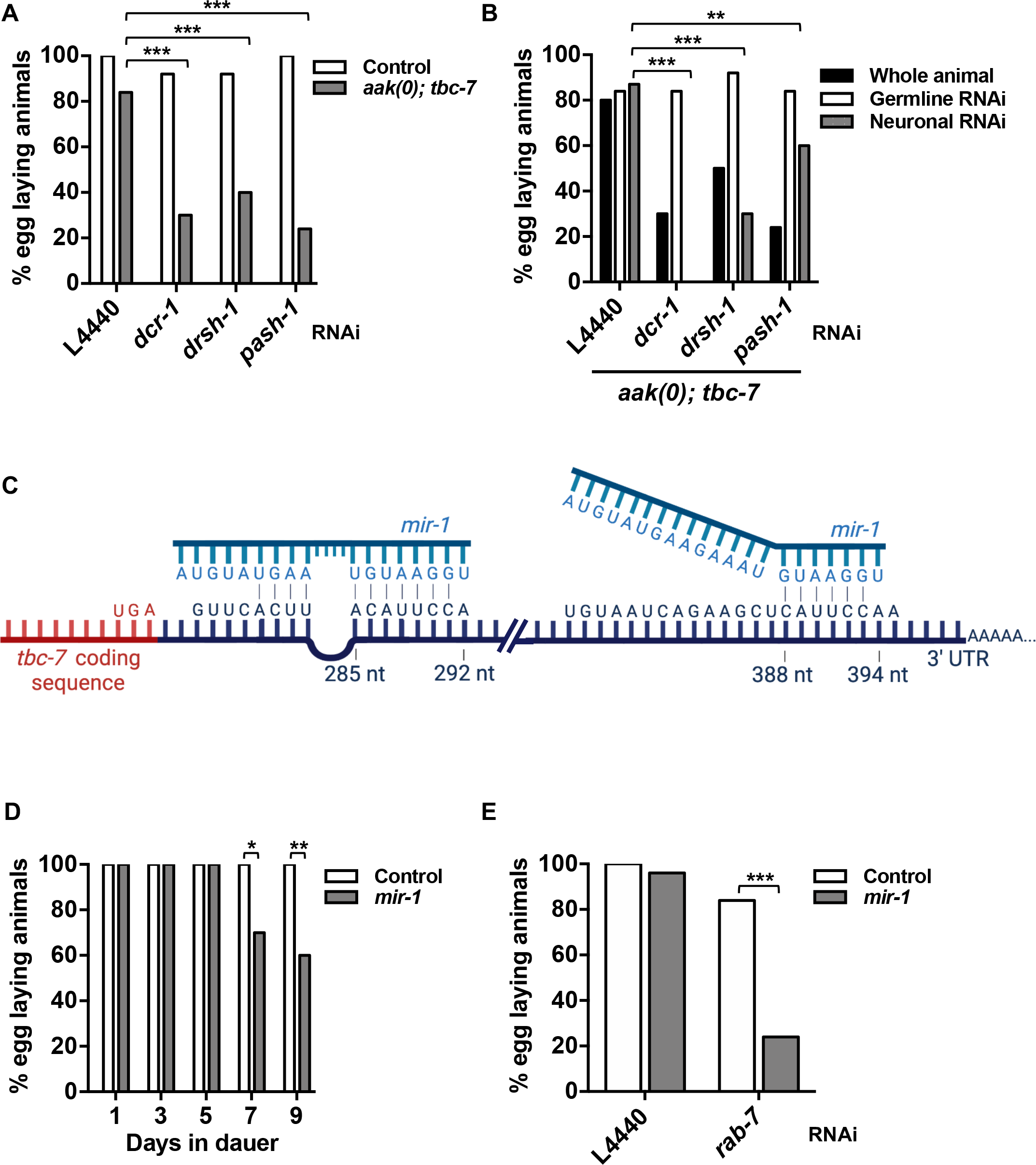
*mir-1* regulates *tbc-7* expression in the neurons to maintain germ cell integrity during the dauer stage. (A) The observed *tbc-7*-associataed suppression of the AMPK germline phenotypes requires the function of miRNA biogenesis components. ***P < 0.0001 when compared to L4440 empty vector in *aak(0); tbc-7* using Marascuilo procedure for % of egg laying animals. Control refers to post-dauer *daf-2(e1370)*. (B) Tissue-specific RNAi experiments were performed to identify the tissues where the miRNA biogenesis components are required for the *tbc-7* suppression of AMPK germline phenotypes. ***P < 0.0001, **P < 0.001 when compared to L4440 empty vector in neuronal RNAi using Marascuilo procedure for % egg laying animals. (C) A model illustrating the binding of *mir-1* to the two predicted seed sequences on the 3’ UTR of *tbc-7*. Lines indicate predicted binding between *mir-1* and *tbc-7* 3’ UTR. (D) *mir-1* mutants exhibit post-dauer sterility after 7 days in the dauer stage. **P < 0.001, *P < 0.01 when compared to control animals that spent the same number of days in the dauer stage using Marascuilo procedure for % egg laying animals. (E) TBC-7 exerts a dosage-dependent effect on the levels of active GTP-bound RAB-7, and *rab-7* RNAi accelerates this process and causes post-dauer sterility after only 1 day in the dauer stage. ***P < 0.0001 when compared to control using Marascuilo procedure for % egg laying animals. All animals carry the *daf-2(e1370)* allele. n = 50.

To further examine the requirement for small RNAs in this process, we employed the same tissue-specific RNAi strategy to determine in which tissues the small RNA function is required in. Surprisingly, disabling the microRNA biogenesis machinery in the germ line did not affect the post-dauer fertility of the *aak(0); tbc-7* mutants, while the loss of the miRNA biogenesis machinery in the neurons reverts the suppression of AMPK germline defects conferred by *tbc-7*, suggesting that miRNA production is not required in the germ line, but rather in the neurons (Fig 6B). Together these data indicate that the suppression of the AMPK mutant post-dauer sterility conferred by blocking *tbc-7* requires the expression of microRNAs in the neurons in order to restore germline quiescence and germ cell integrity during the dauer stage.

To determine if any potential small RNAs could also impinge on *tbc-7* function, we used TargetScanWorm (version 6.2), a program designed to measure the biological relevance between predicted miRNAs and seed sequences. We identified two highly conserved seed sequences for *mir-1* in the 3’ UTR of *tbc-7* (Table S5). *mir-1* is one of 32 conserved microRNAs throughout the Bilateria and is important for many functions including autophagy, muscle development, and sarcomere and mitochondrial integrity [27, 28].

If *mir-1* directly regulates the expression of *tbc-7*, then the loss of *mir-1* during the dauer stage should result in the abnormal upregulation of TBC-7. This increase in TBC-7 activity would further enhance the hydrolysis of RAB-7 GTP into RAB-7 GDP, resulting in the same post-dauer phenotypes seen in *aak(0)* mutants. When *mir-1* mutants were allowed to spend 24 hours in the dauer stage, there were no observable defects in the post-dauer fertility (Fig 6D). However, if the duration of dauer is prolonged to 7 days, *mir-1* mutants exhibited varying degrees of post-dauer sterility, while control animals were unaffected (Fig 6D). Furthermore, when the levels of the TBC-7 target RAB-7 were reduced by RNAi, the *mir-1* mutants exhibited post-dauer sterility after only 24 hours in the dauer stage (Fig 6E). These data suggest that *mir-1* regulates *tbc-7* in a dose-dependent manner. As the animal spends an extended period of time in the dauer stage, the loss of *mir-1* allows for *tbc-7* to be actively expressed. TBC-7 then will continually enhance the hydrolysis of RAB-7 GTP into RAB-7 GDP, decreasing the amount of active RAB-7 available. It is this depletion of active RAB-7 in its GTP-bound state that is responsible for the loss of germ cell integrity, rendering the post-dauer animals sterile (Fig 7).

**Fig 7.**
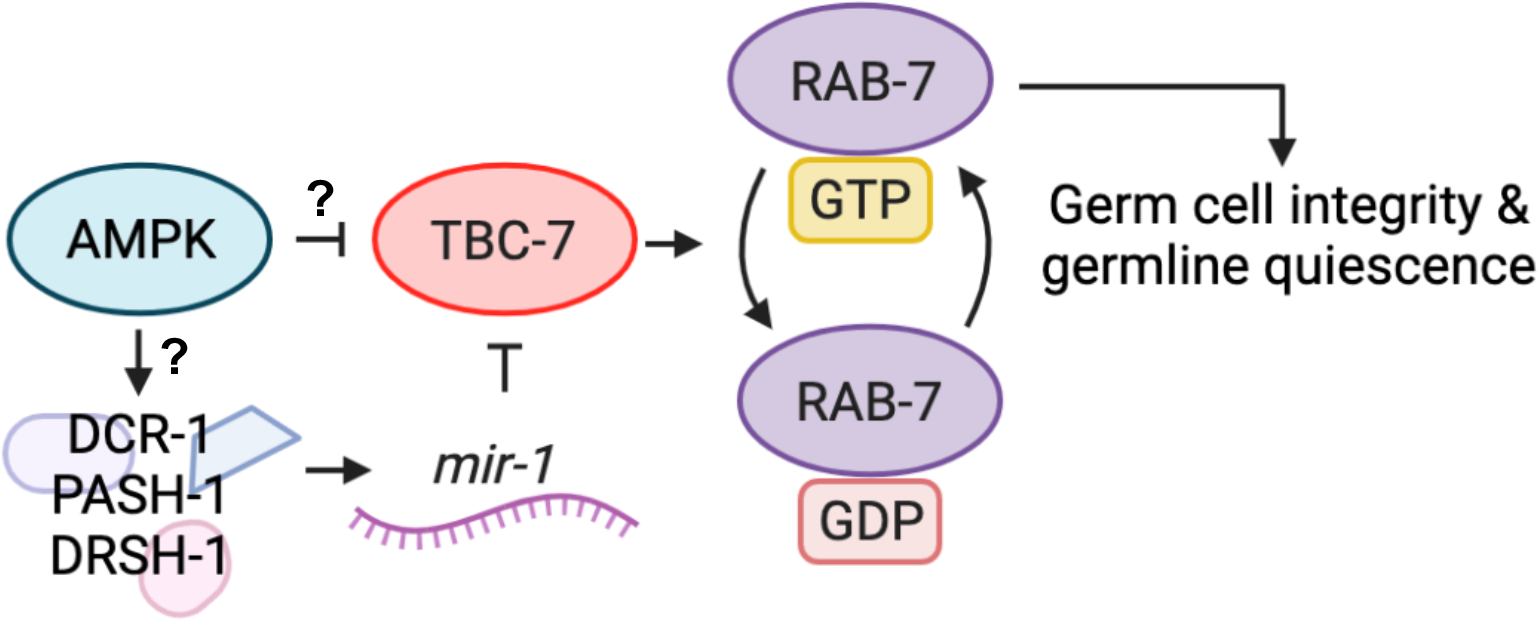
AMPK regulates *rab-7* activity in neurons to establish germline quiescence and preserve germ cell integrity during the dauer stage. When animals enter the dauer stage, AMPK regulates the miRNA biogenesis in the neurons, either directly or indirectly, to generate *mir-1*. *mir-1* is then able to bind to the 3’UTR of *tbc-7* to negatively regulate TBC-7 expression. In parallel, AMPK may also act directly on TBC-7 through phosphorylation to inhibit its activity or destabilize it. Reducing TBC-7 activity allows RAB-7 to remain in its active GTP-bound form to mediate changes in the dauer germ line to preserve germ cell integrity and post-dauer reproductive fitness.

## Discussion

Upon entry into the developmentally arrested dauer stage, AMPK modulates the germline cell cycle, while also triggering changes to preserve germ cell integrity throughout the diapause. This is achieved in part by regulating the chromatin landscape and the associated changes in germline gene expression [10]. In the absence of this master metabolic regulator, animals exhibit germ cell hyperplasia during the dauer stage, while animals that are allowed to recover exhibit highly penetrant sterility as well as various somatic defects. This correlates with an increased abundance and abnormal distribution of both activating and repressive chromatin marks in the germ cells of these mutants. These modifications are likely responsible for the dramatic deviation from the normal expression of germline-specific genes in AMPK mutant dauer larvae and this maladaptive expression program persists into the post-dauer animals [10, 14].

Using genetic analysis to identify suppressors of this reproductive defect, we isolated 7 mutants that corresponded to 5 complementation groups. Two of these groups had multiple alleles and one of these mutations corresponded to a RabGAP protein called TBC-7. TBC-7 null mutants are embryonic lethal, underscoring once again the power of unbiased forward genetic approaches in the identification of highly specific point mutations. Curiously, the EMS induced lesion results in a serine to phenylalanine change within an exon. Genetic analysis indicates that this mutation is hypomorphic and likely produces a RabGAP protein that is unable to reduce the activity of its target.

TBC-7 is important for the proper maintenance of germ cells during this period of quiescence. When *tbc-7* activity is compromised, the defects observed in the chromatin landscape in the germ line of the AMPK mutant dauer larvae are corrected to near wild type levels. This pathway may therefore be one of the major effectors of germline quiescence that occurs during the dauer stage. Curiously, the orthologues of this gene product are important for vesicle formation in neurons, and in *C. elegans* TBC-7 is expressed almost exclusively in neurons, not in the germ cells.

We have previously shown that restoring AMPK signalling within the neurons is sufficient to correct both the germline and the post-dauer somatic defects of AMPK mutants [10]. The neuron-specific expression of *aak-2* did not ameliorate the premature lethality of AMPK mutant dauer larvae [16]. Similarly, although the disruption of *tbc-7* has a remarkable effect on the germline defects of AMPK mutants, loss of *tbc-7* did not improve the premature lethality associated with the metabolic misregulation characteristic of AMPK mutant dauer larvae. Our data therefore suggest that the tissue-specific requirements for regulating germ cell integrity differ from those necessary for dauer survival, and that the neuronal activity of *tbc-7/rab-*7 probably has no significant role in regulating metabolism during the dauer stage.

The nervous system constitutes the perfect early response system to mediate any organismal adaptation to environmental challenges. Since AMPK activity in these cells is sufficient to change gene expression in the germ line, this protein kinase likely acts as a direct sensor for cellular ATP levels to transduce a signal from the neurons to vulnerable cell types dispersed throughout the organism [10, 29].

*tbc-7* is expressed in many neurons in both dauer larvae and adult animals, where it regulates the activity of RAB-7, a small *ras*-like GTPase involved in the positive regulation of the early to late endosome transition, where it localizes to the late endosomal membrane [30, 31]. Late endosomes are involved in recycling cargo back to the plasma membrane, trafficking towards the Golgi body, or fusion with lysosomes to create a lysosome/late endosome hybrid to degrade cargo [30–32]. Given this wide range of functions, the mechanistic details of how RAB-7 may communicate information between the neurons and the germ line remain unclear. Our observations support a role for RAB-7 in the cellular trafficking of a neuron-derived diffusible signal to the germ line to establish germ cell quiescence during the dauer stage, although the nature of this diffusible signal remains enigmatic.

We have recently shown that small RNAs are involved in maintaining germ cell integrity during the dauer stage [10]. Upon the inhibition of endogenous siRNA activity, both the post-dauer fertility and the upregulation of chromatin marks in AMPK mutants are partially corrected. Furthermore, the loss of the dsRNA channel *sid-1* partially restores post-dauer germ cell integrity in the AMPK mutants, implying that the transfer of endogenous siRNAs during the dauer stage is misregulated and maladaptive in the absence of AMPK.

We demonstrate here that disabling microRNA biogenesis components reverts the suppression of AMPK germline defects conferred by *tbc-7*, while suppressing the biogenesis of the ALG-3/4 and ERGO-1 26G classes of siRNAs has no observable phenotype. Furthermore, our tissue-specific RNAi studies show that microRNA biogenesis is only required in the neurons, suggesting that this small RNA signal originates in these cells. Our findings indicate that the aberrant activity of siRNAs during the dauer stage may be harmful to the animal and that AMPK could initiate the neuron-specific production of microRNAs required to preserve germline integrity during the dauer stage. Our data suggest that AMPK engages the miRNA pathway both upstream of *tbc-7* through the action of *mir-1* and downstream of *tbc-7* to mediate the effects of activated RAB-7.

AMPK must act upstream of miRNA biogenesis and corroborates previous findings by Han et al and Jose et al who have provided evidence that small RNAs produced in the neurons can exert cell non-autonomous effects [33, 34]. It is however still unknown how AMPK is able to regulate small RNA production or function. DRSH-1, PASH-1, and DCR-1 have multiple medium stringency AMPK phosphorylation motifs, so it is possible that AMPK directly phosphorylates the microRNA biogenesis machinery to ensure that microRNAs are produced in a timely manner. Alternatively, it is possible that microRNA production is regulated independently of AMPK/LBK1 signalling and these protein kinases impinge on components downstream of miRNA biogenesis to mediate these changes in a permissive manner. Further work will be required to discern between these possibilities.

By analysing the microRNA seed sequences present within the 3’UTR of *tbc-7*, we were able to identify and characterise how one microRNA, *mir-1*, is critical for the maintenance of germ cell integrity. Animals that lack *mir-1* show the same germline defects as AMPK mutants, although these phenotypes are only visible after these mutants spend a longer period of time in the dauer stage. These germline defects are further accentuated upon the compromise of *rab-7* expression, suggesting that in the *mir-1* mutants *tbc-7* continually suppresses the activation of *rab-7*. This is also the case in other organisms where *mir-1* has been shown to be a regulator of synaptic function at neuromuscular junctions [27]. This genetic data thus places *mir-1* in a linear pathway as a negative regulator of *tbc-7* expression, and therefore, indirectly, as a positive regulator of *rab-7* activity. At the onset of dauer, AMPK activation might ultimately affect the formation of GTP-bound RAB-7, which in turn could act as the major effector in the neurons enabling communication with the germ cells to ensure germ cell integrity.

Although we have described how *mir-1* contributes to the regulation of *tbc-7* in the neurons in response to AMPK activation, the kinetics of the *mir-1* inhibition are not sufficient to account for the timing of the changes in the germ line following the initiation of dauer formation. When *rab-7* is compromised using RNAi in the *aak(0); tbc-7* mutant, the animals exhibit highly penetrant post-dauer sterility after spending 48 hours in the dauer stage. On the contrary, *mir-1* mutants must spend a considerably longer period of time in this stage before they exhibit post-dauer sterility, suggesting that *mir-1* may not be the sole regulator of *tbc-7* expression. TBC-7 possesses a high stringency AMPK phosphorylation motif on Ser115. During wild type dauer formation, AMPK could phosphorylate this key serine residue on TBC-7 to negatively affect the function or stability of TBC-7 in addition to *mir-1* regulation. Our hypomorphic *tbc-7(rr166)* mutation could affect the function or stability of TBC-7 such that it is no longer functional, thus mimicking the effects of both *mir-1* regulation and AMPK phosphorylation causing the suppression of AMPK germline defects. These possibilities could be examined by quantifying the levels of TBC-7 during the onset of dauer in control animals and AMPK mutants. If the abundance of TBC-7 changes, this data could indicate that phosphorylation by AMPK may affect its stability and target it towards the proteasome. The AMPK-mediated phosphorylation of TBC-7 would then account for the major pathway destabilizing the RabGAP as animals form dauer larvae, while the *mir-1* regulation of the *tbc-7* transcript would act redundantly and in parallel to ensure that TBC-7 is incapable of blocking RAB-7 activation during the dauer stage.

In summary, we have characterised an AMPK-dependent *mir-1/tbc-7/rab-7* pathway that acts cell non-autonomously in the neurons to establish quiescence in the germ line during periods of energetic stress. As the animals enter the dauer stage, *mir-1* is produced downstream of DCR-1 and DRSH-1/PASH-1, which could potentially be activated by an AMPK-dependent phosphorylation. *mir-1* then binds to the 3’UTR of *tbc-7* to negatively regulate its expression, thus allowing RAB-7 to remain in its GTP-bound active form and to perform its function. The miRNA pathway is then engaged once again to mediate the RAB-7 mediated process in maintaining germ cell integrity and germline quiescence during the dauer stage (Fig 7). Although we have identified this *mir-1*/*tbc-7*/*rab-7* pathway, we have yet to identify which particular function of *rab-7* is responsible for transmitting this dauer signal from the neurons to the germ line. Our findings provide mechanistic details of the regulation of the *mir-1*/*tbc-7*/*rab-7* pathway occurring in the neurons to alter the chromatin landscape and enable the subsequent changes in gene expression required to establish germline quiescence during periods of energetic stress.

## Materials and Methods

### *C. elegans* strains and maintenance

All *C. elegans* strains were maintained at 15°C and according to standard protocols unless otherwise indicated [35]. The strains used in this study is listed in the following table:

| Strain | Genotype |
| --- | --- |
| CB1370 | *daf-2(e1370) III* |
| MR1000 | *daf-2(e1370) aak-1(tm1944) III; aak-2(ok523) X* |
| MR1963 | *daf-2(e1370) aak-1(tm1944) III; tbc-7(rr166) aak-2(ok523) X* |
| MR2267 | *daf-2(e1370) aak-1(tm1944) III; tbc-7(rr267) aak-2(ok523) X* |
| MR2265 | *daf-2(e1370) aak-1(tm1944) III; rr249 V; aak-2(ok523) X* |
| MR2204 | *daf-2(e1370) aak-1(tm1944) III; rr256 aak-2(ok523) X* |
| MR2205 | *daf-2(e1370) aak-1(tm1944) III; rr266 aak-2(ok523) X* |
| MR2269 | *daf-2(e1370) aak-1(tm1944) III; rr289 aak-2(ok523) X* |
| MR2270 | *daf-2(e1370) aak-1(tm1944) III; rr261 aak-2(ok523) X* |
| MR2268 | *daf-2(e1370) aak-1(tm1944) III; rr268 IV; aak-2(ok523) X* |
| DCL569 | *mkcSi13 [sun-1p::rde-1::sun-1 3'UTR + unc-119(+)] II; rde-1(mkc36) V* |
| MR2292 | *mkcSi13 [sun-1p::rde-1::sun-1 3'UTR + unc-119(+)] II; daf-2(e1370) aak-1(tm1944) III; rde-1(mkc36) V; aak-2(ok524) X* |
| MR2359 | *daf-2(e1370) aak-1 (tm1944) III; rde-1(mkc36) V; aak-2(ok524) X* |
| MR2283 | *mkcSi13 [sun-1p::rde-1::sun-1 3'UTR + unc-119(+)] II; daf-2(e1370) aak-1(tm1944) III; rde-1(mkc36) V; tbc-7(rr166) aak-2(ok524) X* |
| MR2308 | *daf-2(e1370) aak-1 (tm1944) III; rde-1(mkc36) V; tbc-7(rr166) aak-2(ok524) X* |
| VC576 | *mir-1(gk276) I* |
| MR2289 | *mir-1(gk276) I; daf-2(e1370) III* |
| MR2324 | *tbc-7(rr166) X* |
| FX31259 | *tbc-7(tm10766)/tmC30(tmls1247) X* |

Transgenic lines and compound mutants were created in the laboratory using standard molecular genetics approaches. All mutant strains isolated following mutagenesis were backcrossed at least 5 times prior to characterization and whole-genome sequencing.

### Genetic suppressor screen

Animals were mutagenized as described elsewhere with some modifications [35]. P_0_ L4 AMPK null (*aak(0)*) mutants were treated with 0.05 mM EMS. F_1_ animals were isolated to obtain F_2_ candidates that were homozygous for the EMS-generated mutation. F_2_ animals were switched to restrictive temperature to induce dauer formation and allowed to spend 48 hours in this stage before they were switched to permissive temperature to trigger dauer recovery. Five F_2_ post-dauer progeny from each F_1_ parent were isolated, allowed to form dauer then were subjected to 1% SDS in order to eliminate dauer-defective mutants. Potential candidates were further screened for post-dauer fertility to identify suppressors of the germline defects typical of AMPK mutants. In total, 8 independent alleles were isolated from approximately 7000 haploid mutagenized genomes.

### Complementation analysis

Complementation analysis was conducted as previously described following injection of *myo-2p*::GFP and heat shock to induce males. Males with the *myo-2p*::GFP transgene were crossed with mutant adult hermaphrodites, using pharyngeal GFP as a marker for cross progeny. The F_1_ heterozygotes were transited through the dauer stage and post-dauer fertility was assessed as a measure of complementation.

### Dauer recovery assay

Post-dauer fertility and somatic phenotypes assay was conducted as described elsewhere [10].

### DAPI staining and germ cell quantification

DAPI staining and germ cell quantification of the dauer larvae was conducted as described elsewhere [10].

### Dauer survival assay

Dauer survival assay was conducted as described elsewhere [16].

### Western blot analysis

*C. elegans* dauer larvae and post-dauer adults were diluted in lysis buffer (50 mM Hepes [pH 7.5], 150 mM NaCl, 10% glycerol, 1% Triton X-100, 1.5 mM MgCl_2_, 1 mM EDTA, and protease inhibitors cocktail [Halt^TM^, 100X, Thermo Scientific]) and lysed by multiple cycles of freeze-boiling. Protein concentrations were determined using a NanoDrop 2000C spectrophotometer (Thermo Fisher Scientific, Waltham, MA, USA). Nitrocellulose membranes were incubated with primary antibodies: rabbit anti-H3K4me3 (1:1000 dilution, Abcam, ab8580), rabbit anti-H3K9me3 (1:1000 dilution, Cell Signalling Technology, 9754S), and mouse anti-α-tubulin (1:2000 dilution, Developmental Studies Hybridoma Bank, AA4.3). Proteins were visualized using horseradish peroxidase-conjugated goat anti-rabbit secondary antibody conjugate (1:2000 dilution, Bio-Rad laboratories, #1706515) or horseradish peroxidase-conjugated goat anti-mouse secondary antibody conjugate (1:2000 dilution, Bio-Rad laboratories, #1706516). Membranes were incubated in Clarity^TM^ western ECL substrate (Bio-Rad laboratories, #1705061). Relative protein abundance of α-tubulin and histone marks were determined using Image J. Membranes were imaged using MicroChemi (DNR Bio-Imaging Systems) and E-Gel Imager GelCapture Acquisition System (ThermoFisher).

### Immunostaining and quantification

*C. elegans* dauer larvae and post-dauer adult gonads were dissected, fixed, and stained as described elsewhere [36]. Extruded gonads were incubated with rabbit anti-H3K4me3 (1:500 dilution, Abcam, ab8580) or rabbit anti-H3K9me3 (1:500 dilution, Cell Signalling Technology, 9754S). Secondary antibodies were Alexa-Fluor-488–coupled goat anti-rabbit (1:500; Life Technologies, Carlsbad, CA, USA). Gonads were counterstained with DAPI (0.1 μg/mL dilution, Roche Diagnostics, 10236276001). Microscopy was performed as described elsewhere [37]. Ratios for the fluorescence intensity across the germ line were determined using Image J.

### RNA interference by feeding

RNA interference was conducted as described elsewhere based on standard feeding protocols [10, 38].

### Generation of tissue-specific RNAi strains

All tissue-specific RNAi strains contain a *rde-1(mkc36)* mutation. *rde-1(mkc36)* contains a 67 bp insertion and a 4 bp deletion that creates three premature stop codons [20]. *rde-1(mkc36)* mutants do not respond to exogenous RNAi by feeding or injection [20]. A germline-specific RNAi strain was generated by expressing a single-copy of *rde-1* in the germ line driven by the *sun-1* promoter inserted on chromosome II using MosSCI [20]. To generate neuron-specific RNAi strains, *rde-1* and *sid-1* were expressed exclusively in the neurons of *rde-1(mkc36)* animals using the *rgef-1* promoter [39]. Neurons of *C. elegans* exhibit weak penetrance through feeding, thus extra copies of neuronal *sid-1* potentially serves as a dsRNA sink, allowing for a robust RNAi phenotype through feeding [21]. Tissue-specific RNAi strains do not exhibit any developmental defects when grown in replete conditions, when passed through the dauer stage on standard NGM plates, or when grown on L4440 empty vector control compared to their non-*rde* counterpart.

### Prediction of *mir-1* seed sequence

*mir-1* was identified as a potential regulator of *tbc-7* through TargetScanWorm release 6.2. Two highly conserved *mir-1* seed sequences were identified in the 3’UTR of *tbc-7*. The P_CT_ and conserved branch length are measures of the biological relevance of the predicted miRNA and target interaction, with greater values being more likely to have detectable biological function [40, 41].

## Acknowledgements

We thank the members of the Roy and Zetka laboratories for their thoughtful discussions and comments on the manuscript. We acknowledge the *Caenorhabditis elegans* Genetics Center and the National BioResource Project: *Caenorhabditis elegans* Shigen for *C. elegans* strains. This work was supported by the Canadian Institutes for Health Research (CIHR).

## Supporting information

**S1 Fig.**
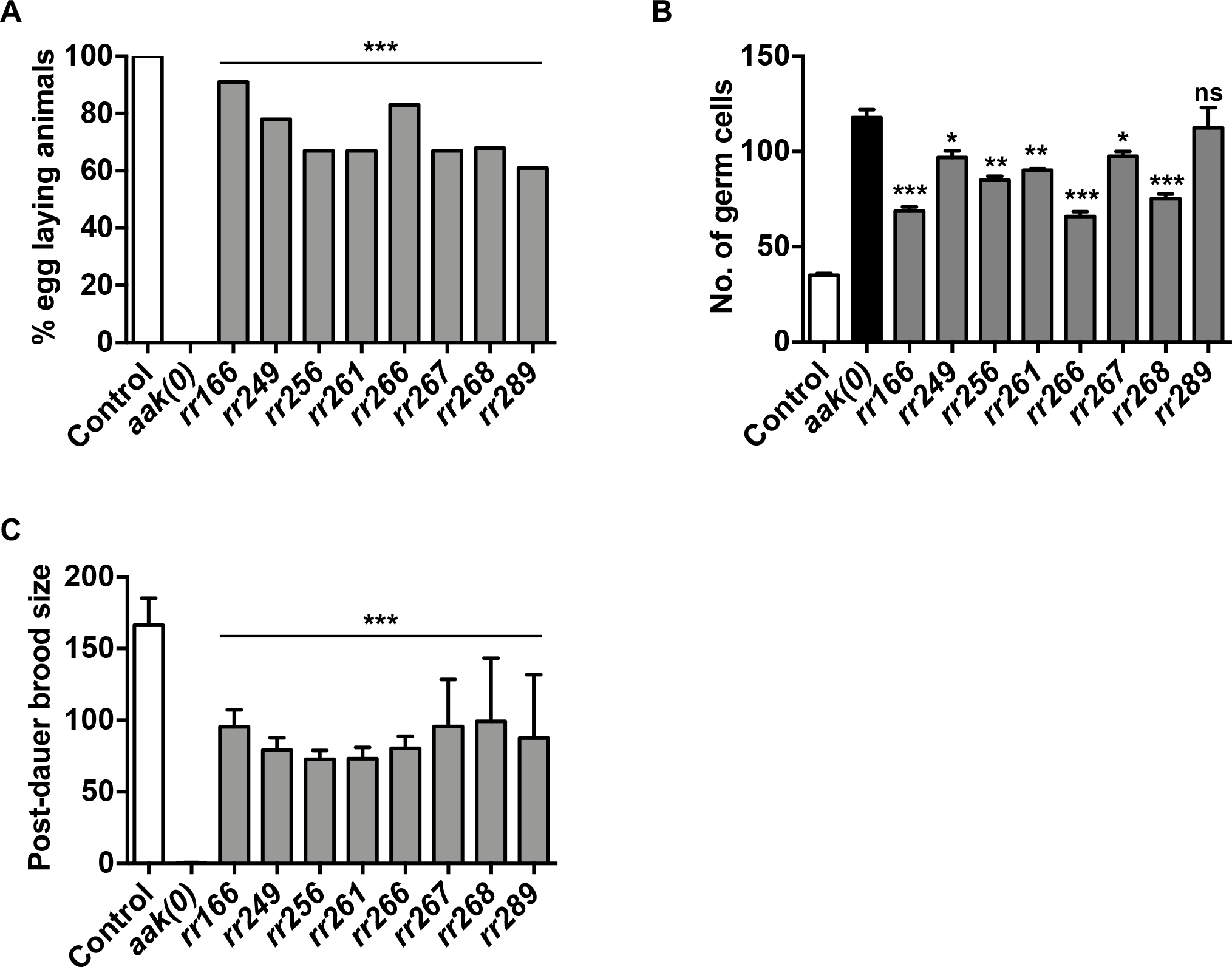
Mutants partially suppress the dauer germline hyperplasia and post-dauer sterility associated with loss of AMPK. (A-C) Mutants isolated from an EMS suppressor screen partially suppress the (A) post-dauer sterility, (B) dauer germline hyperplasia, and (C) brood size defects associated with a lack of AMPK signalling. ***P < 0.0001, **P < 0.001, *P < 0.05 when compared to *aak(0)* using ordinary one-way ANOVA for mean brood size and mean germ cells. ***P < 0.0001 when compared to *aak(0)* using Marascuilo procedure for % egg laying animals and somatic phenotypes. All animals isolated from the screen include *daf-2; aak(0)* in the background. n = 50.

**S2 Fig.**
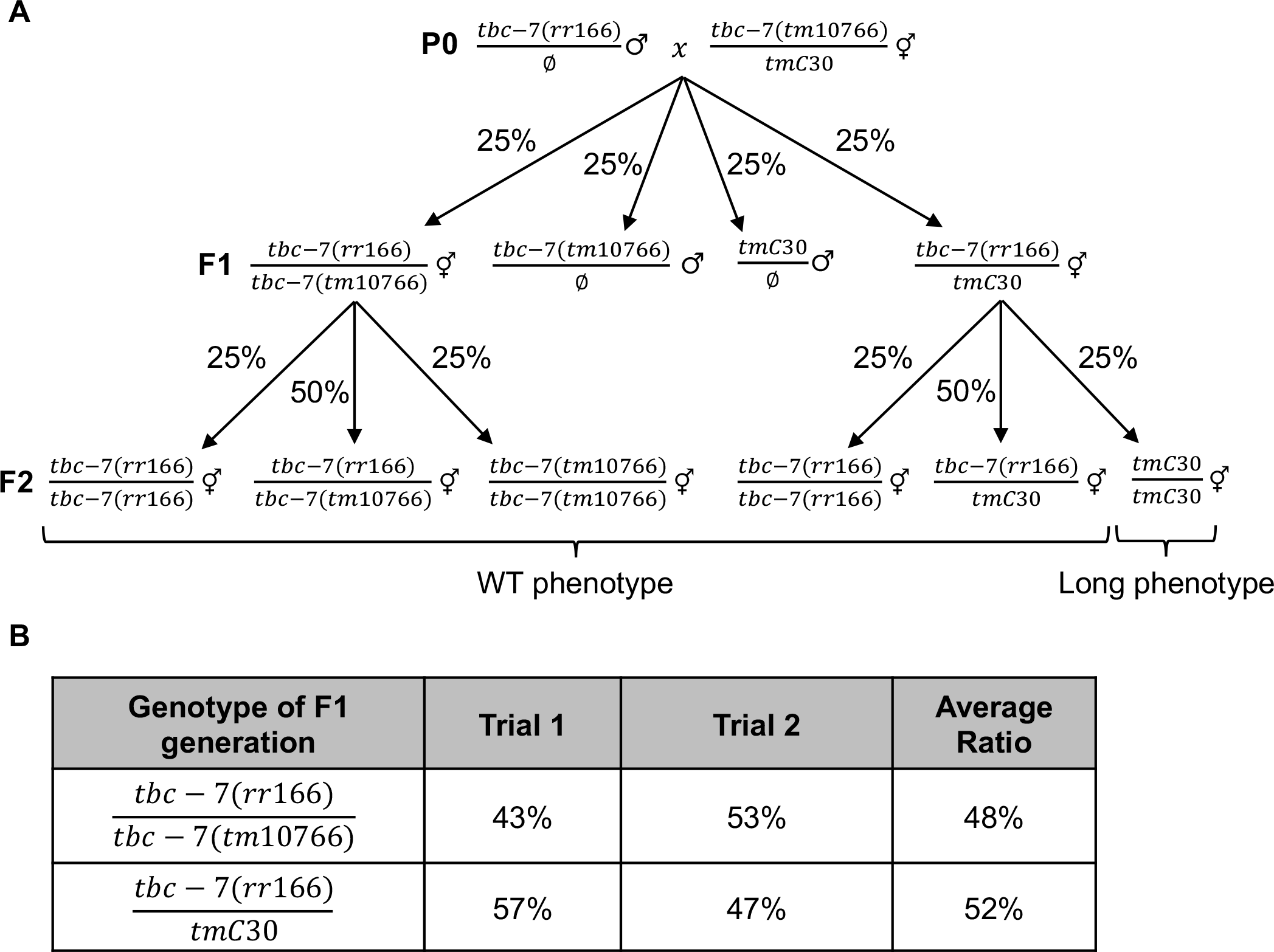
*tbc-7(rr166)* is a hypomorphic allele. (A) Schematic showing the cross with *tbc-7(rr166)* with *tbc-7(tm10766)*, which contains a 21,846 bp deletion that completely removes *tbc-7*. Homozygous *tbc-7(tm10766)* is lethal and is balanced with *tmC30*, which is a balancer that has a recessive Long (Lon) phenotype. To confirm if *tbc-7(rr166)* is a hypomorphic allele, *tbc-7(rr166)* was crossed with *tbc-7(tm10766)* and the F_2_ cross progeny were examined in order to identify F_1_ heterozygous *tbc-7(rr166)/tbc-7(tm10766)* hermaphrodites. F_1_ heterozygous *tbc-7(rr166)/tbc-7(tm10766)* hermaphrodites should yield no Lon phenotype progeny, while F_1_ heterozygous *tbc-7(rr166)/tmC30* should yield 25% Lon phenotype progeny. (B) Table showing the percentage of F_1_ genotypes over two independent crosses. The F_2_ progeny of every F_1_ was assessed. F_1_ genotypes assessed per cross n ≥ 150.

**S3 Fig.**
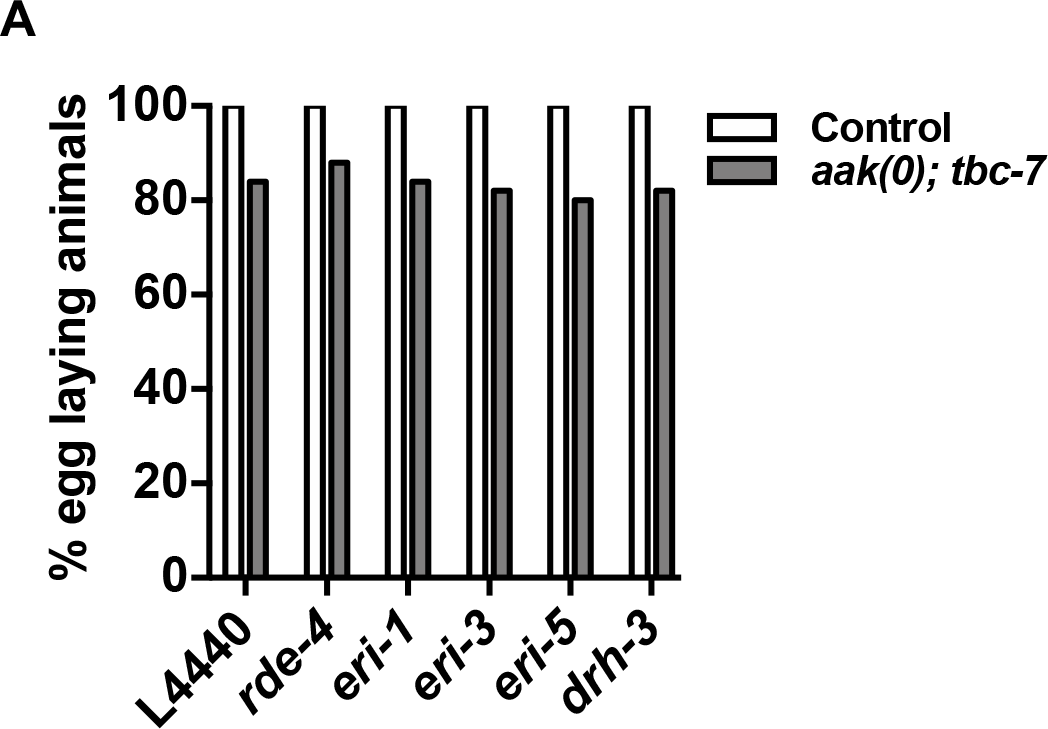
*tbc-7* is not regulated by small RNAs produced from the ERGO-1 26G and ALG-3/4 26G classes. (A) The compromise of ERGO-1 and ALG-3/4 class biogenesis factors have no quantifiable effect on the post-dauer fertility of *tbc-7*-suppressed mutants. All values are ns when compared to control. ns when compared to L4440 empty vector using Marascuilo procedure for % egg laying animals. All animals carry the *daf-2(e1370)* allele. n = 50.

---

**S1 Table.**
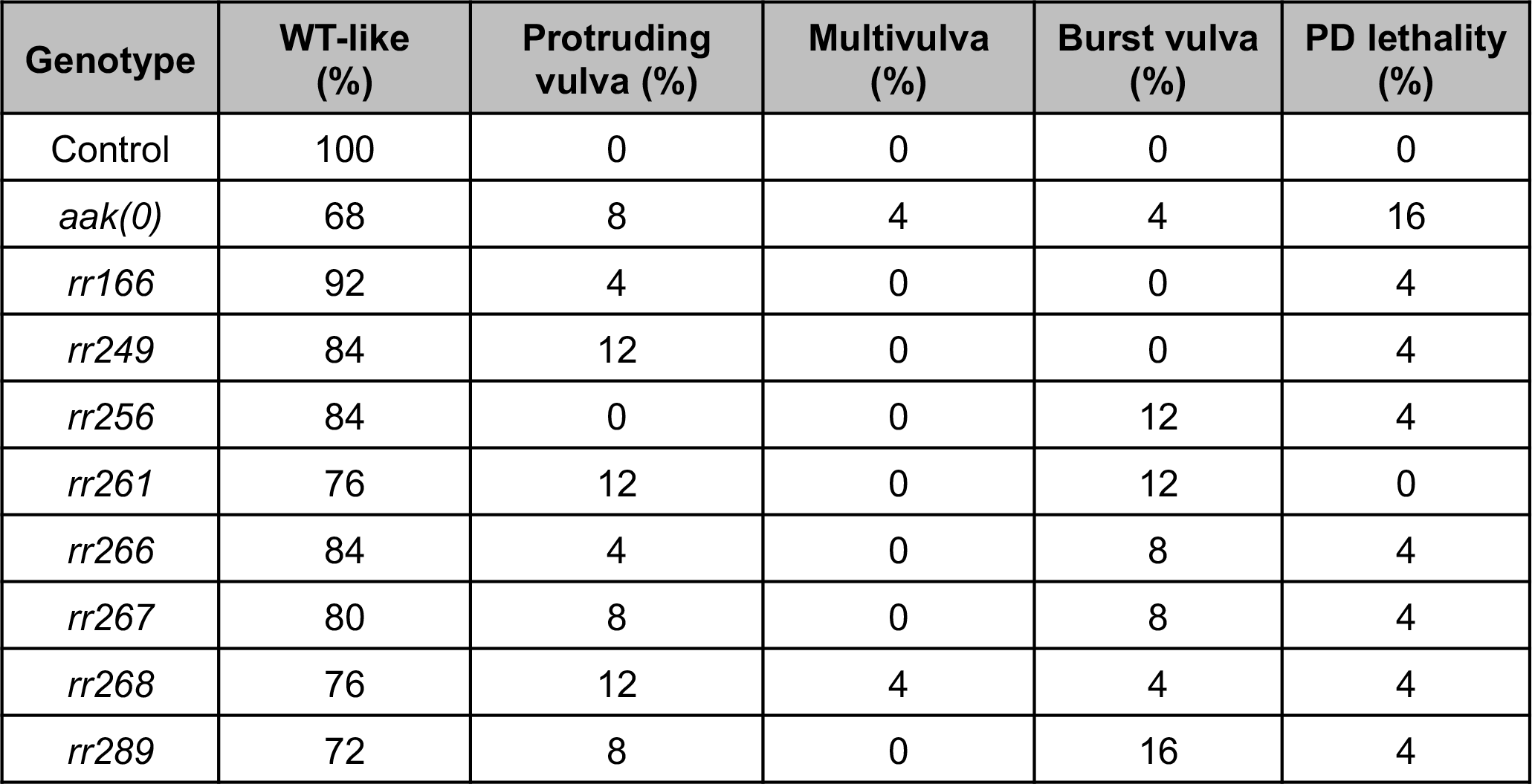
Mutants isolated from an EMS suppressor screen partially suppress the post-dauer somatic defects of AMPK mutants. Mutants were allowed to transit through the dauer stage and recover. The post-dauer somatic defects were assessed seven days after the recovery period. All animals carry the *daf-2(e1370)* allele. Mutants isolated from the EMS screen are *aak(0)*. n = 50.

| Genotype | WT-like (%) | Protruding vulva (%) | Multivulva (%) | Burst vulva (%) | PD lethality (%) |
| --- | --- | --- | --- | --- | --- |
| Control | 100 | 0 | 0 | 0 | 0 |
| *aak(0)* | 68 | 8 | 4 | 4 | 16 |
| *rr166* | 92 | 4 | 0 | 0 | 4 |
| *rr249* | 84 | 12 | 0 | 0 | 4 |
| *rr256* | 84 | 0 | 0 | 12 | 4 |
| *rr261* | 76 | 12 | 0 | 12 | 0 |
| *rr266* | 84 | 4 | 0 | 8 | 4 |
| *rr267* | 80 | 8 | 0 | 8 | 4 |
| *rr268* | 76 | 12 | 4 | 4 | 4 |
| *rr289* | 72 | 8 | 0 | 16 | 4 |

**S2 Table.**
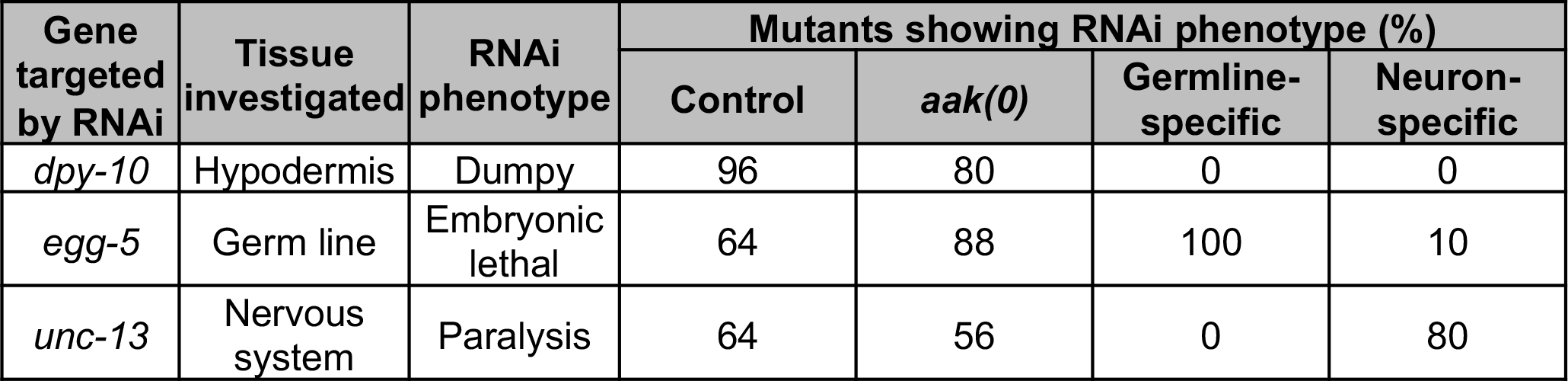
*aak(0)* mutants with tissue-specific expression of *rde-1* exhibit tissue-specific RNAi phenotypes. To confirm that *aak(0)* mutants with tissue-specific expression of *rde-1* exhibit tissue-specific RNAi phenotypes, mutants were fed dsRNA against *dpy-10* (hypodermis), *egg-5* (germ line), or *unc-13* (neurons) and the phenotypes were scored. Only mutants expressing *rde-1* in the same tissues targeted by the dsRNA should exhibit the RNAi phenotype. All mutants are *daf-2; aak(0)*.

**S3 Table.** Post-dauer fertility after RNAi treatment against *rab* genes. Control and *aak(0); tbc-7* mutants were fed dsRNA against all known and predicted *rab* genes then allowed to transit through the dauer stage. L4440 empty vector was used as a control. n = 50.

|  | % egg laying animals |  |
| --- | --- | --- |
| RNAi | Control | *aak(0); tbc-7* |
| L4440 | 100 | 84 |
| *rab-1* | 0 | 0 |
| *rab-2* | 60 | 40 |
| *rab-3* | 100 | 64 |
| *rab-5* | 24 | 0 |
| *rab-6.1* | 95 | 44 |
| *rab-6.2* | 86 | 84 |
| *rab-7* | 84 | 12 |
| *rab-8* | 83 | 64 |
| *rab-10* | 84 | 20 |
| *rab-11.1* | 68 | 40 |
| *rab-11.2* | 7 | 8 |
| *rab-14* | 84 | 44 |
| *rab-21* | 85 | 88 |
| *rab-27* | 100 | 84 |
| *rab-28* | 80 | 92 |
| *rab-35* | 100 | 60 |
| *rab-37* | 96 | 64 |
| *rab-39* | 96 | 52 |
| R07B1.12 | 100 | 84 |
| 4R79.2 | 100 | 56 |
| K02E10.1 | 100 | 52 |
| F11A5.3 | 96 | 56 |
| F11A5.4 | 100 | 56 |
| C33D12.6 | 100 | 84 |
| C33E6.22 | 100 | 84 |

**S4 Table.** *aak(0); tbc-7* mutants with tissue-specific expression of *rde-1* exhibit tissue-specific RNAi phenotypes. To confirm that *aak(0); tbc-7* mutants with tissue-specific expression of *rde-1* exhibit tissue-specific RNAi phenotypes, mutants were fed dsRNA against *dpy-10* (hypodermis), *egg-5* (germ line), or *unc-13* (neurons) and the phenotypes were scored. Only mutants expressing *rde-1* in the same tissues targeted by the dsRNA should exhibit the RNAi phenotype. All mutants are *daf-2; aak(0); tbc-7*.

| Gene targeted by RNAi | Tissue investigated | RNAi phenotype | Mutants showing RNAi phenotype (%) |  |  |  |
| --- | --- | --- | --- | --- | --- | --- |
|  |  |  | Control | *aak(0); tbc-7* | Germline-specific | Neuron-specific |
| *dpy-10* | Hypodermis | Dumpy | 96 | 84 | 0 | 0 |
| *egg-5* | Germ line | Embryonic lethal | 64 | 72 | 100 | 0 |
| *unc-13* | Nervous system | Paralysis | 64 | 68 | 0 | 80 |

**S5 Table.**
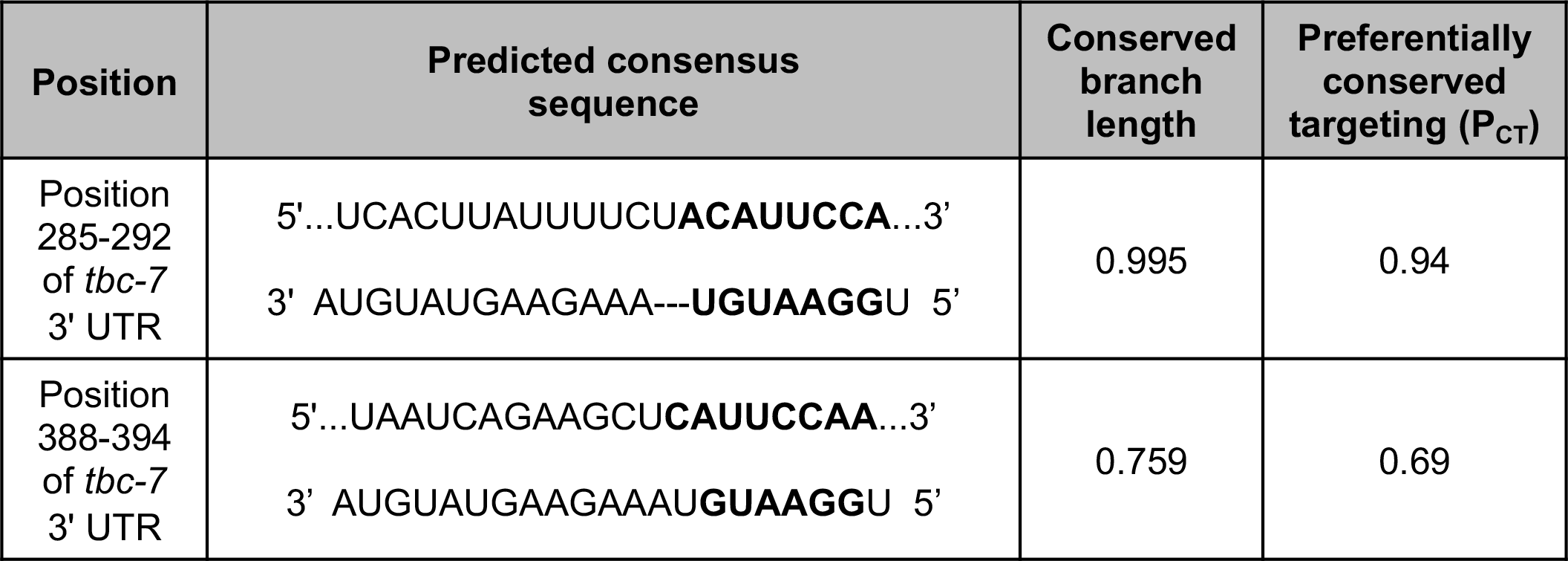
Predicted *mir-1* seed sequences in the 3’ UTR of *tbc-7*. Two highly conserved *mir-1* seed sequences (bold) were identified in the 3’ UTR of *tbc-7* (TargetScanWorm release 6.2), suggesting that *mir-1* directly regulates *tbc-7* expression. The P_CT_ and conserved branch length are measures of the biological relevance of the predicted miRNA and target interaction, with greater values being more likely to have detectable biological nction [40, 41].

